# rG4-seq 2.0: enhanced transcriptome-wide RNA G-quadruplex structure sequencing for low RNA input samples

**DOI:** 10.1101/2022.02.10.479665

**Authors:** Jieyu Zhao, Eugene Yui-Ching Chow, Pui Yan Yeung, Qiangfeng Cliff Zhang, Ting-Fung Chan, Chun Kit Kwok

## Abstract

RNA G-quadruplexes (rG4s) are non-canonical structural motifs that have diverse functional and regulatory roles such as transcription termination, alternative splicing, mRNA localization and stabilization and translational process. We recently developed RNA G-quadruplex structure sequencing (rG4-seq) technique and discovered many rG4s in both eukaryotic and prokaryotic transcriptomes. However, rG4-seq suffers from complicated gel purification step and limited PCR product yield and thus requires a high RNA input amount, limiting its applications for physiologically or clinically relevant studies. In this study, we have developed rG4-seq 2.0 by introducing a new ssDNA adapter containing deoxyuridine in the library preparation to enhance the library quality with no gel purification step, less PCR amplification cycles and higher yield of PCR products. We demonstrate that rG4-seq 2.0 produced high quality cDNA libraries that supported reliable and reproducible rG4 identification at varying RNA inputs (as low as 10 ng amount of RNA). rG4-seq 2.0 also improved the rG4-seq calling outcome and nucleotide bias in rG4 detection persistent in rG4-seq 1.0. Our new method can improve the identification and study of rG4s in low abundance transcripts, and our findings can provide insights to optimize cDNA library preparation in other related methods.

## INTRODUCTION

RNA structure plays an important role in biological function and regulation including but not limited to transcriptional and translational processes, RNA splicing, polyadenylation, mRNA localization and stabilization [1–6]. During the last decade, a number of methods were developed to study RNA structures by combining enzymatic and chemical probing with next generation sequencing (NGS) technique [7–16], yielding comprehensive *in vitro* and *in vivo* structural map of RNA on a transcriptome-wide level, and providing new insights to RNA biology [7–16]. One unique RNA secondary structure, RNA G-quadruplex, which is composed of G-quartets connected by loop nucleotides, is of emerging importance in both chemistry and biology fields, and have been associated with diseases such as cancers and neurological disorders [6, 17–21]. In 2016, we introduced RNA G-quadruplex sequencing (rG4-seq) [22] for the identification of *in vitro* rG4 on a transcriptome-wide scale that can facilitate further exploration and study of *in vivo* rG4 structure and function. Recently, we have refined a few steps of the rG4-seq experimental procedures [23] and also developed a bioinformatics pipeline known as rG4-seeker [24]. To date, rG4-seq has been successfully applied to study the rG4s in human [22], plant [25], bacteria [26], and plasmodium [27].

It is noted that 5’ adapter ligation step is one of the critical steps in rG4-seq and several other cDNA library preparation methods [22, 28–30]. Previously, we invented a hairpin DNA adapter to hybridize with the incoming cDNA and perform the ssDNA ligation mediated by T4 DNA ligase [31]. Over the years, this identical or highly similar strategy has been widely applied in the ligation step of new library preparation methods [23, 32–36]. The major limitation of such hairpin DNA adapter is that the secondary structure of the adapter may interfere with the base-pairing of forward primer with the cDNA template. Furthermore, the complicated gel purification step after the 5’ adapter ligation also lengthens experimental processing time and increases the samples loss. Both factors result in lower PCR amplification efficiency, requiring more PCR cycles (or input RNA) to obtain sufficient double-stranded DNA (dsDNA) for sequencing (Figure 1A).

**Figure 1.**
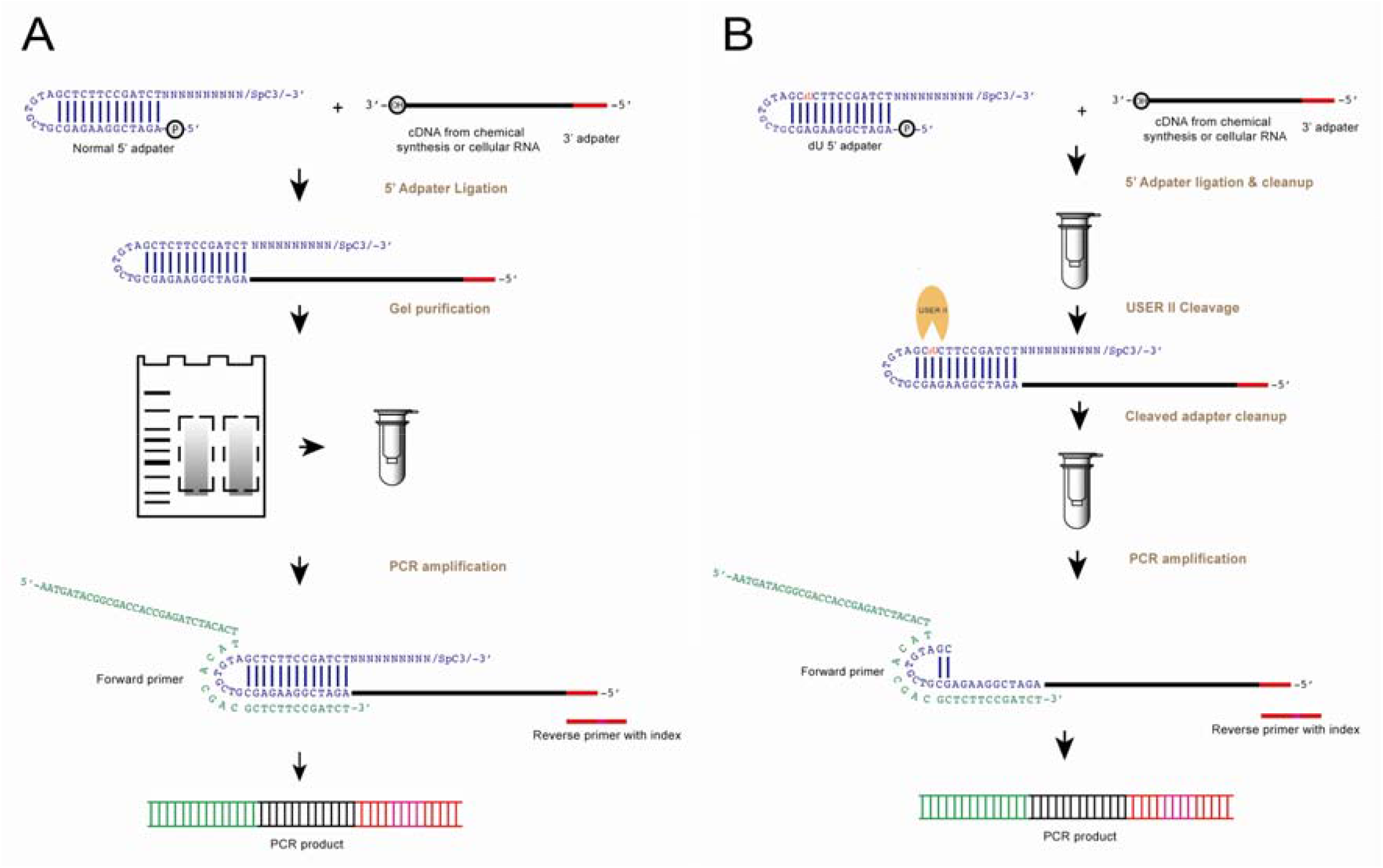
Experimental flowchart of the 5’ adapter ligation and dU cleavage. The sequences of the oligonucleotides used can be found in Table S1. The experimental pipelines that use **(A)** normal 5’ adapter and **(B)** 5’ dU adapter are shown (blue). In A), the normal 5’ adapter ligates to the cDNA (black) with 3’adapter sequence (red) followed by gel purification before performing PCR amplification. The 6nt index region in the reverse PCR primer is in pink. In B), dU adapter is used for the ligation, and after the column purification, the enzymatic cleavage step is performed with cleavage enzyme (USER II, orange pacman) followed by column clean-up before the PCR amplification. USER II enzyme cleaves the 5’ adapter at the dU position as indicated in red in the 5’ dU adapter. The dU adapter contains the same secondary structure as the normal 5’ adapter, and carries a 5’ phosphate at its 5’ end to ligate to incoming cDNA with a 3’hydroxyl, and a C3 spacer group at its 3’ end to block and avoid self-oligomerization. The 10 random nucleotide (N10) serve as a template for the hybridization of any incoming cDNA.

In this study, we have introduced a new 5’ DNA adapter that contains a deoxyuridine (5’ dU adapter) and removed the gel purification step (Figure 1B) in the library preparation of rG4-seq, which we referred as rG4-seq 2.0. Using model oligonucleotides, we successfully applied rG4-seq 2.0 to varying polyA-RNA inputs (500 ng, 100 ng, 30 ng, and 10 ng) and showed that the data are of high quality, reproducibility and accuracy for transcriptome mapping and rG4 calling. Compared to old method [23], rG4-seq 2.0 yielded higher PCR products with less PCR cycles and improved rG4 calling outcome by mitigating a significant nucleotide bias in rG4-induced RTS site identification persistent in the old method. We reported for the first time the *in vitro* rG4 map of HEK293T cells, and discussed the relationships among RNA amount inputs, read coverage and transcript abundance for rG4 mapping. Furthermore, we propose a standard recommendation for better judgement of RNA amount inputs and sequencing depth for specific rG4 study based on the target abundance in the transcriptome.

## MATERIALS AND METHODS

### Preparation of DNA oligonucleotides

All the DNA oligonucleotides used in this work were synthesized and purchased from IDT. The detailed sequences were listed in the Table S1. The oligos were dissolved and diluted to desired volume with nuclease-free water accordingly. N40+22 nt DNA template was PAGE purified before use.

### USER II cleavage-The effect of reaction time

A 20 μl reaction mixture was made of 2 μl of 10 μM of either normal or dU adapter (1 μM final), 2 μl of 10X CutSmart® buffer (1X final) and 2 μl of 1U/μl USER® II enzyme (M5505S, NEB) (0.1U/μl final) and 14 μl nuclease-free water. The reaction mixture was incubated at 37°C for 0, 1, 2, 5 and 15 minutes. For test of reaction time over 15 min, same 20 μl reaction mixture was prepared and incubated at 37°C for 15, 30, 60 and 120 minutes. At each time point, 5 μl of mixture were aliquoted out and added with 5 μl 2X formamide (FA) dye for quenching. The samples were subjected to gel electrophoresis afterwards (See below).

### USER II cleavage-The effect of enzyme concentration

The reaction mixture was made of 1 μl of 10 μM dU adapter (1 μM final), 1 μl of 10X CutSmart® buffer (1X final), 0, 0.5, 1 or 2 μl of 1U/μl USER® II enzyme (M5505S, NEB) and nuclease-free water to made up to 10 μl. The reaction mixture was incubated at 37°C for 15 minutes. 10 μl 2X FA dye was added to each sample. Gel electrophoresis was then performed (See below).

### USER II cleavage-The effect of uracil position

The reaction mixture was composed of 1μl of 5’ dU adapter / dU Loop18 adapter / dU Loop20 adapter /dU stem27 adapter / dU stem32 adapter / dU stem34 adapter (1 μM final), 1 μl of 10X CutSmart® buffer, 7 μl nuclease-free water and 1 μl of USER® II enzyme. The reaction was incubated at 37°C for 15 minutes. After that, 10 μl 2X FA dye was added to the sample. Gel electrophoresis was then performed (See below).

### 5’ dU adapter ligation-The effect of DNA:adapter ratio on the ligation efficiency

The ligation reaction was carried out using Quick Ligation™ kit (NEB, M2200). A 10 μl mixture consist of 1 μl of 1 μM N40+22nt DNA (0.1 μM final), 1 μl of 5’ dU adapter with varying concentration (0.1, 0.25, 0.5 or 1 μM final), 2 μl of 50% PEG 6000, 1 μl of Quick ligase in 1X Quick ligation buffer. The reaction condition was 2 hours at 37°C and 10 minutes at 95°C for inactivation followed by adding with 10 μl 2X FA dye. The samples were subjected to gel electrophoresis (See below).

### Purification steps - The effect of column purification on the USER II cleavage after ligation

5’ adapter ligation was performed with 5 pmol DNA and 50 pmol dU adapter input in the 20 μl volume reaction. After 2 hours incubation at 37 °C and 10 minutes at 95°C, for reactions without column purification, 5 μl of reaction was taken out followed by adding 5 μl nuclease-free and 10 μl 2X FA dye for reference. Another 5 μl of reaction was directly added with 4 μl nuclease-free water and 1 μl of USER® II enzyme and incubated at 37°C for 15 minutes. For reaction with column purification, 10 μl of ligated product was added to 40 μl nuclease-free water before performing the column purification with RNA Clean & Concentrator™ (Zymo Research, R1016). After that, 10 μl nuclease-free water was eluted. 5 μl was taken out and added with 5 μl nuclease-free water followed by 10 μl 2X FA dye as reference. The other 5 μl reaction was mixed with 3 μl nuclease-free water, 1 μl CutSmart® buffer and 1 μl of USER® II enzyme, and then incubated at 37°C for 15 minutes and quenched with 2X FA dye. The samples were subjected to gel electrophoresis (See below).

To test recovery rate of column purification on the ligated product, 20 μl 106-nt mimic ligated product (0.25 μM final) was prepared. 10 μl was taken out for column purification with RNA Clean & Concentrator™ (Zymo Research, R1016) and eluted with 10 μl of nuclease-free water. The other 10 μl reaction was used for reference. each reaction was quenched with 2X FA dye. The samples were subjected to gel electrophoresis (See below).

### Purification steps - The effect of column purification on the cleaved adapters removal after USER II cleavage

To perform excess adapter removal after USER II cleavage, 20 μl of 5’ adapter ligation and column purification after ligation were stated above. Eluted the sample with 16 μl nuclease-free water followed by 2 μl CutSmart® buffer and 2 μl USER® II treatment. After 15 min incubation at 37°C, take out 10 μl and add 40 μl nuclease-free for second column purification (Zymo Research, R1016) followed by elution with 10 μl nuclease-free water. Another 10 μl sample was saved for reference control. For size references, nuclease-free water was added to replace the dU adapter or N40+22 DNA template in preparation of 5’adapter ligation reaction. Each reaction was quenched with 10 μl 2X FA dye and gel electrophoresis was performed. (See below).

### Denaturing polyacrylamide gel electrophoresis (PAGE)

Samples were heated at 95°C for 3 minutes before loading into the gel. Each sample (2 μl) was then loaded into the pre-heated (300 V for 20 minutes) 8.3 M urea-containing 10% polyacrylamide-urea denaturing gel. The gel electrophoresis was performed at 300 V for 25 minutes in 1X TBE buffer.

To visualize the nucleic acids, SYBR Gold (Thermo, S11494) was used to stain the gel for 5 minutes after electrophoresis. The gel was then soaked into the staining solution for 5 minutes and then directly scanned by Bio-rad ChemiDoc™ Touch Imaging System and the gel image was analyzed.

### PCR amplification

100 ng polyA-enriched RNAs from HEK 293T cell were used to generate cDNA for PCR amplification using normal adapter following rG4-seq [23] and 5’ dU adapter (See library preparation method below). The PCR reaction included 4 μl cDNA template (from either normal or 5’ dU adapter sample), 0.5 μl of 10 μM PCR forward primer (0.5 μM final), 0.5 μl of 10 μM PCR reverse primer (0.5 μM final), 5 μl of 2X KAPA HiFi HotStart Ready Mix (KAPA Biosystems, KK2602).

The PCR program was stated below: 95 °C, 3mins; (98 °C, 20 seconds; 68 °C, 15 seconds; 72 °C, 40 seconds), 72 °C, 1.5 minutes. The PCR amplifications were conducted in 4 different cycles, i.e. 8, 11, 14, 17. After the amplification, the samples were subjected to 2% agarose gel electrophoresis at 120V for 60 minutes. The gel image was obtained for further analysis.

### Data processing

All the gel bands were background-corrected and quantified by the Image Lab™ 6.0.1 software.

A: Uncleaved adapter

B: Cleaved adapter

C: N40+22 nt DNA template band

D: Ligated product without USER II treatment

E: Ligated product after USER II treatment

F: 106-nt mimic ligated adapter band without column purification as reference

G: 106-nt mimic ligated adapter band with column purification

H: Cleaved adapters before column purification as reference

I: Cleaved adapter after column purification

#### Cleavage rate

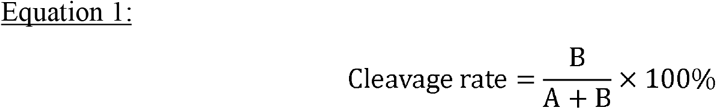

#### Reduction rate of Ligated product

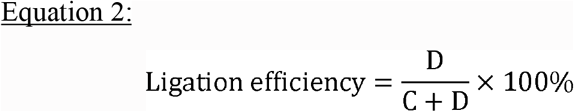

#### 5’ adapter ligation efficiency

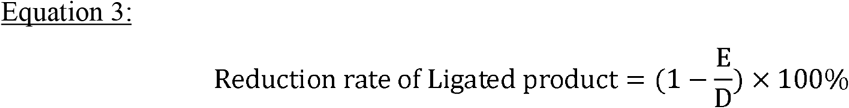

#### Recovery rate of column purification

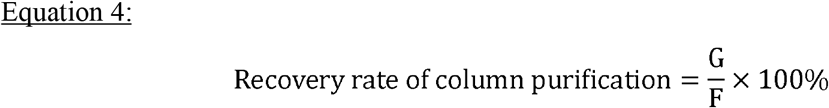

#### Cleaved adapters removal rate

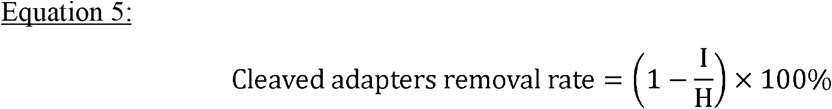

### High throughput library preparation and sequencing data analysis

Authenticated HEK293T cells without mycoplasma contamination were grown in DMEM (Thermo Scientific, 10569044) media with 10% heat inactivated FBS (Thermo Scientific, 10270106) and 1X antibiotic antimycotic (Thermo Scientific, 15240062) at 37 °C with 5% CO2. Total RNA was extracted from the pelleted cells using RNeasy Plus Mini Kit (Qiagen, 74136).

PolyA RNAs were purified using the Poly(A) PuristMAG Kit (Thermo Scientific, AM1922). After 2 rounds of selection, The corresponding amount of polyA RNA were incubated with 1X fragmentation buffer at 95 °C for 1 minute to generate ~300 nt of RNA fragment followed by column purification (Zymo Research, R1016) as described before [23]. Next, 3’ dephosphorylation was conducted at 37 °C for 30 minutes including 7 μl sample after fragmentation, 1 μl rSAP enzyme (NEB, M0371L) and 1 μl PNK enzyme (NEB, M0201S) in 1X T4 PNK buffer. Then 3’-adapter ligation was performed at 25°C for 1 hour by adding 1μl 3’ adapter (with 1:10 molar ratio of RNA to 3’adapter), 7 μl PEG 8000 (17.5% final) and 1 μl of T4 RNA ligase 2 KQ (NEB, M0373L) in 1X T4 RNA ligase buffer. After 3’adapter ligation, added 1 μl RecJf (NEB, M0331) and 1 μl 5’deadenylase (NEB, M0331) into the 20 μl ligation reaction for incubation with 30 minutes at 30°C and 3 minutes at 65°C to inactivate the enzymes followed by column purification (Zymo Research, R1016). Before the reverse transcription (RT), each sample was then equally separated into 2 reactions. Each reaction includes 12 μl of sample, 1 μl reverse primer (with 1:2.5 molar ratio of RNA to reverse primer), 6 μl Li^+^ or K^+^ RT buffer [23]. RT process was described in previous study [23]. After RT, cDNA was purified for further 5’adapter ligation. At this step, 1 μl 5’ dU adapter (with 1:10 molar ratio of cDNA to 5’ adapter) was mixed with 7 μl eluted cDNA sample and then was heated at 95 °C for 3 minutes, 60 °C for 1 minute and 25 °C for 2 minutes. After the incubation, 10 μl 2X Quick T4 ligase buffer and 2 μl Quick T4 DNA ligase (NEB, M2200L) were added to the reaction and incubate at 37 °C for overnight. The ligated products (≥200 nt) were then purified by RNA Clean & Concentrator (Zymo Research, R1016). Subsequent steps including dU adapter cleavage with CutSmart® buffer and column purification were stated in detail above. After the PCR cycle test, real PCR reaction of each sample including 8 μl purified ligated product after dU cleavage, 1 μl 10 μM forward primer, 1 μl 10 μM reverse primer with different 6-nt index sequence (barcode) and 10 μl 2X KAPA HiFi HotStart ReadyMix (KAPA Biosystems, KK2602), was performed using 95 °C, 3 minutes, appropriate cycles of each temperature step (98 °C, 20 seconds; 65 °C, 15 seconds; 72 °C, 40 seconds) based on the PCR test result. Then PCR products with loading dye were purified with 2% agarose gel, and sizes 150-400 bp were sliced and extracted (Zymo research, D4008). Following steps were performed similar as previously described [23]. The rG4-seq 1.0 libraries with 100 ng polyA-enriched RNA input were performed based on previous method [23]. Library quantification was obtained by qPCR (KAPA, KK4824). Each replicate of quantified samples was then pooled and subjected to next-generation sequencing on the Illumina Hiseq System.

### Sequencing data processing and bioinformatic analysis

UMI barcode in raw sequencing reads were first extracted using UMI-tools [37]. Sequencing reads were then adapter-trimmed and quality-trimmed using cutadapt [38]. Trimmed reads were aligned to GRCh38/hg38 human reference genome using STAR aligner [39]. Uniquely aligned reads were deduplicated based on UMI information using UMI-tools[37]. Deduplicated and uniquely aligned reads were subjected to rG4-seeker pipeline [24] to identify RTS sites and rG4 motifs at default settings. Ensembl 97 gene annotation [40] was used to define the transcriptomic regions. Transcript abundances (in TPM) were quantified using Kallisto [41] employing all aligned reads, and were then summed to obtain gene abundances. Downstream analysis to intersect RTS sites detected in respective rG4-seq libraries and to match RTS sites with parent gene abundances were conducted with Python, where two RTS sites with <5nt distance on genomic coordinates were considered overlapping. Consensus lists of rG4s detected across rG4-seq 2.0 and rG4-seq 2.0/1.0 libraries were obtained with iterative runs of bedtools intersect [42] and using 500 ng-input rG4-seq 2.0 library as basis. Union lists of rG4s detected across rG4-seq 2.0 and rG4-seq 1.0 libraries were obtained using bedtools merge [42], where adjacent/overlapping rG4 motifs on the same strand from different libraries were merged.

## RESULTS AND DISCUSSION

### USER II Cleavage

#### Effect of reaction time on normal and 5’ dU adapter cleavage by USER II

5’ adapter ligation is an essential step before PCR amplification in the cDNA library preparation of rG4-seq. We hypothesize that the 5’ adapter with strong secondary structure may affect with downstream processes such as PCR amplification efficiency (Figure 1). To test this, based on our previously developed normal 5’adapter (Figure 1A), we first designed a modified 44-nt dU adapter with a dU nucleotide at the 24^th^ nucleotide position from 5’ end to replace the T nucleotide (Figure 1B). To process the dU adapter, Thermolabile USER II enzyme (USER II) was used, which creates the abasic site and further cleaves out the phosphodiester backbone to produce 2 DNA fragments, with the size of 23 nt and 21 nt, from the 5’ dU adapter. To explore the appropriate reaction time for dU cleavage, we tested and compared the cleavage rate with different reaction time on both dU and normal adapter.

We performed the USER II treatment at different time points (0, 1, 2, 5, 15 minutes) on dU adapter and 0 and 15 min for normal adapter as negative controls, and ran them on denaturing PAGE gel to determine if the 5’ dU adapter (44 nt) was cleaved into shorter fragments (23 nt and 21 nt) by USER II. Compared to normal 5’ adapter (Figure 2, lanes 1-2), dU adapter showed varying degrees of cleavage with increasing incubation time (Figure 2, lanes 3-7). The cleavage rate reached over 96% in 15 min reaction time (Figure 2, lane 7), while there was no cleavage observed at 0 min for dU adapter (Figure 2, lane 3). As expected, no cleavage was observed on normal adapter in both 0 min and 15 min (Figure 2, lanes 1 and 2). We also tested dU cleavage with longer time including 30, 60 and 120 minutes, and found there was no significant difference on the cleavage efficiency compared to 15 minutes (Figure S1A). Therefore, we found that USER II could specifically cleave dU adapter at the dU position with a highest cleavage rate in 15 minutes of reaction time.

**Figure 2.**
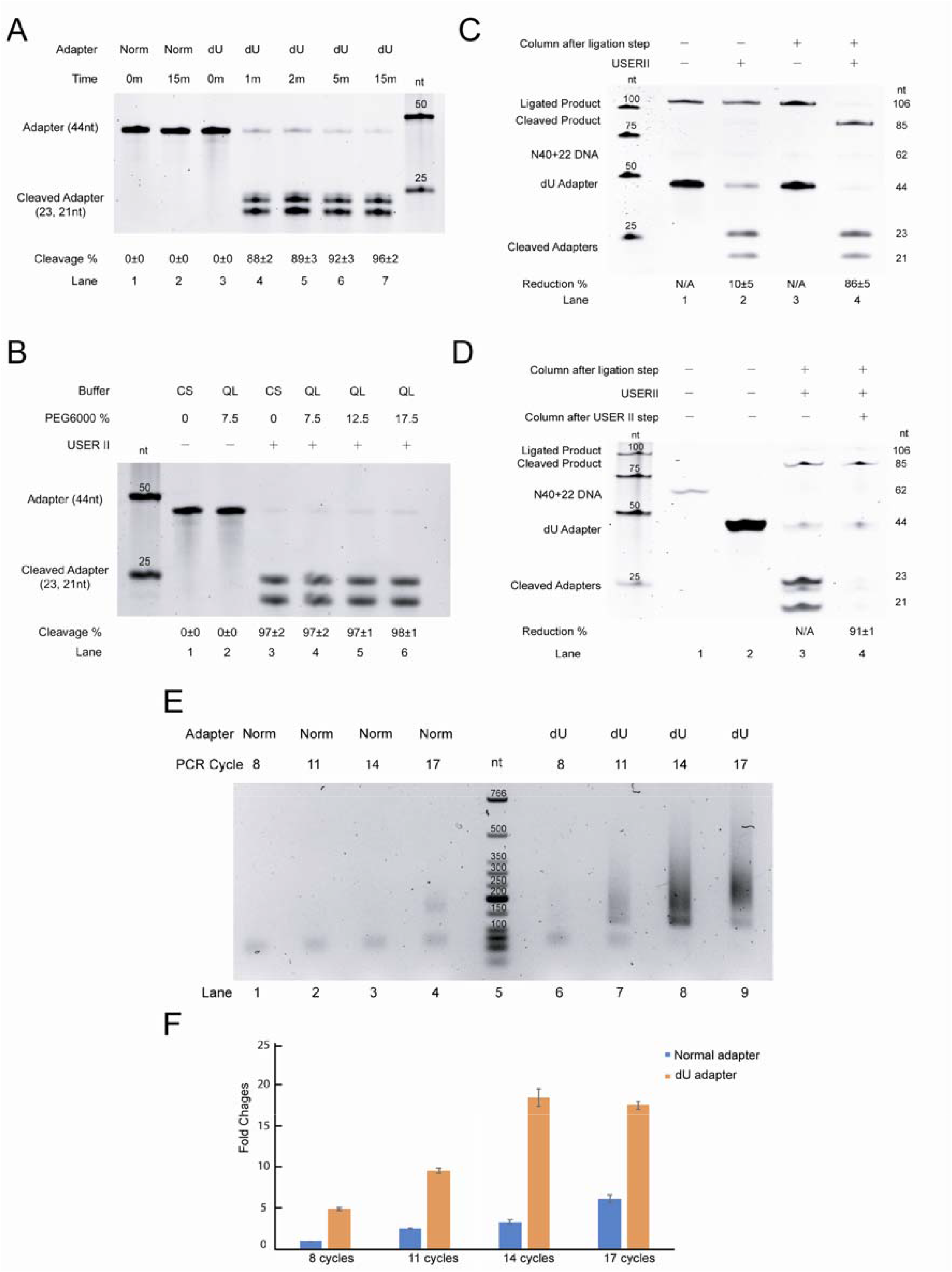
Optimization of experimental procedures. **(A)** The effect of reaction time on the normal and 5’ dU adapter cleavage. The figure shows the cleavage of both normal (lane 1-2) and dU adapter (lane 3-7) with USER II at different reaction time. **(B)** The effect of reaction buffer and different percentage of PEG 6000 on the dU 5’ adapter cleavage. The figure shows the dU cleavage under CutSmart (CS) buffer or Quick Ligase (QL) Buffer for dU adapter (Lane 1-3) and with the percentage of PEG 6000 ranging from 7.5%-17.5% (Lane 4-6). Equation 1 is used for calculating the cleavage rate (See Methods). **(C)** The effect of column purification on the USER II cleavage after ligation step The 106-nt ligated product bands without USER II cleavage before and after column purification (Lanes 1 and 3) are defined as reference for calculation. Equation 3 is used for calculating the reduction rate of ligated product. (See Methods). **(D)** The effect of column purification on cleaved adapters removal after USER II treatment. The cleaved adapters generated from USER II treatment (Lane 3) are used as the reference to calculate the reduction rate of the cleaved adapters after column purification followed by USER II step (Lane 4). The N40+22 DNA only (62 nt in Lane 1) and dU adapter only (44 nt in Lane 2) are used size references. Equation 5 is used for calculating the reduction rate of cleaved adapters (See Methods). **(E, F)** The effect of normal adapter and dU adapter on the PCR amplification. Agarose gel image of PCR result is showed in panel E. PCR products of library generated with 8, 11, 13 and 17 PCR cycles using normal adapter (Norm) and 5’ dU adapter. Comparison of relative quantity of PCR product is showed in panel F. 8 cycles of PCR product with normal adapter is defined as 1. The DNA size marker indicates the fragment size. The errors showed are standard deviation. nt=nucleotide; n=3.

#### Effect of reaction buffer on normal adapter and 5’ dU adapter cleavage

In the workflow (Figure 1), the 5’ dU adapter cleavage step is performed after the 5’adapter ligation in the cDNA library preparation, e.g. in our rG4-seq method [23]. In order to study whether the cleavage efficiency of USER II enzyme is compatible with normal reaction conditions used in the 5’ adapter ligation step, we have assessed two different factors including reaction buffer and PEG 6000 concentration (Figure S1B).

We first compared the cleavage efficiency of USER II enzyme in both CutSmart (CS) (recommended by vendor) and quick ligation (QL) buffer (also used in 5’ adapter ligation). No cleavage was observed regardless reaction buffer type without USER II addition (Figure S1B, lanes 1 and 2), while 97% cleavage rates were obtained after USER II treatment for both CS and QL buffer (Figure S1B, lanes 3 and 4), suggesting that the cleavage efficiency of USER II enzyme was not affected by reaction buffer. Although the commercial QL buffer includes 7.5% PEG 6000 originally, QL buffer got a cleavage rate as high as CS buffer (Figure S1B, lanes 3 and 4). We next tested the effect of different PEG 6000 concentration with QL buffer on the dU adapter reduction rate. Our result showed that from 7.5%-17.5% PEG 6000 concentration, the adapter cleavage rates were all around 97-98% (Figure S1B, lanes 4-6), which were highly similar with the one under CS buffer (Figure S1B, lane 3).

In short, we showed that USER II could work normally with QL buffer and different concentration of PEG 6000 condition.

#### Effect of USER II concetration on 5’ dU adapter cleavage

Besides reaction time and buffer conditions, we have also investigated the effect of USER II concentrations ranged from 0.05U/μl-0.2U/μl on the cleavage efficiency. Five minutes of reaction time was chosen in this test as 5’ dU adapter could be mostly cleaved by USER II enzyme in 15 minutes (Figure 2, lane 7). Our result showed that the cleavage efficiency was 89% under 0.05 U/μl of USER II concentration (Figure S1C, lane 2), while both 0.1 and 0.2 U/μl of USER II concentration leaded to 91% of cleavage rate (Figure S1C, lanes 3 and 4), which showed similar result in Figure 2 (Lane 6). Thus, 0.1U/μl of USER II concentration (Figure S1C, lane 3) is efficient for dU cleavage compared to higher concentration (Figure S1C, lane 4).

#### Effect of dU position on adapter cleavage

dU position at the 24^th^ of nucleotide from 5’ end is designed to increase the base paring ability by introducing cleavage. Besides, we designed two other 5’ dU adapters (dU Loop18 and dU Loop20 adapters, see Table S1 and Figure S2A) with dU at the 18^th^ nt and 20^th^ nt position from 5’end of the adapter to identify whether dU position affects the efficiency of USER II cleavage. Interestingly, our result showed that the cleavage rates of dU Loop18 adapter and dU Loop20 adapter were 37% and 68% (Figure S2A, lanes 2 and 4), which were much lower than the 5’ dU adapter (dU at the 24^th^ position) designed originally (Figure 2), revealing that dU in the single-stranded loop might decrease the USER II cleavage rate compared to dU in the double-stranded DNA (ds DNA). To further verify this, we have also designed three 5’ adapters with dU at 27^th^, 32^th^ and 34^th^ positions (dU Stem27, dU Stem32 and dU Stem34 adapters, see Table 1 and Figure S2B) from 5’ end to identify their cleavage efficiency under 15 minutes of reaction time (Figure S2B). The cleavage rates of dU Stem27, dU Stem32 and dU stem34 adapters by USER II enzyme were 90%, 91% and 84%, respectively (Figure S2B, lanes 2, 4 and 6). In this way, USER II displayed higher efficiency in dsDNA than that of ssDNAs (Figure S2A, lanes 2 and 4). Therefore, we concluded that dU at 24^th^ position of 5’ adapter works well with USER II cleavage.

### 5’ dU Adapter Ligation

To assess the efficiency of 5’ ligation process, a 66-nt nucleotides consist of 40 randomized nucleotides and 22-nt 3’adapter sequence [23] in the 3’end (N40+22, see Table S1), which has been designed to mimic cDNA product after 3’ ligation and reverse transcription in the real library, and ligated to the 44-nt 5’ dU adapter to produce a 106-nt ligated product with the help of T4 DNA ligase. As we have optimized three factors including cDNA: adapter ratio, PEG 6000 concentration and PEG type with normal adapter in previous study [23], here we only examined the effect of dU adapter concentration on the 5’ligation efficiency with other reaction conditions remain unchanged. Four different ratios of cDNA to adapter, ranging from 1:1 to 1:10 were utilized in the 5’ ligation reaction under 17.5% PEG 6000. From the gel image (Figure S3), the ligation yield increased with the reducing DNA: adapter ratio and with 1:5 and 1:10 of DNA:adapter ratios, the ligation efficiency reached 95% (Figure S3, lanes 3 and 4). Overall, we found that 1: 5 or 1:10 of DNA:adapter ratio could get the highest 5’ ligation efficiency with dU adapter.

### Purification Steps Before and After USER II Cleavage

#### The effect of column purification on the ligated product cleavage after ligation

To investigate whether the T4 DNA ligase in the ligation reaction could affect the efficiency of dU cleavage of ligated product after USER II treatment, we performed column purification after the 5’ ligation process to remove those unused substances in the ligation reaction. Even though we showed that the ligation reaction buffer and PEG 6000 concentration had no effect on the dU adapter cleavage (Figure S1B), but whether the whole ligation condition will affect the USER II cleavage rate is still unknown, so here we identified if the 5’ ligation condition affects the following USER II cleavage efficiency with or without ligation cleanup step.

In this study, the entire 106-nt ligated product, including template 44-nt dU adapter at 5’ end and 62-nt cDNA, could be cleaved into 2 fragments (85 nt and 21 nt) by USER II enzyme at dU position (Figure 2C), therefore we evaluated the reduction rate of ligated product with and without column purification to verify if USER II cleavage is compatible with real ligation step. We found that column purification could help USER II cleave up to 86±5% of ligated product (Figure 2C, lane 4) compared to 10±5% of cleavage rate (Figure 2C, lane 2) without column purification after USER II treatment. The 106-nt ligated products without USER II cleavage (Figure 2C, lanes 1 and 3) were reference controls for calculation. As multiple bands showed in the gel image, making it complex to calculate the recovery rate of column purification. Thus, we used a 106-nt oligo to mimic the ligated product (Table 1) and showed the recovery rate of column purification was 92±1% (Figure S4, lane 2), indicating that more than 90% product could be survived during purification. Hence, our result showed that even in QL buffer condition, USER II cleavage cannot compatible with the 5’ ligation condition due to complexity of reaction environment, and a simple column purification after 5’ligation could help to increase the efficiency of USER II cleavage on the ligated product.

#### The effect of column purification on cleaved adapters removal after USER II cleavage

To eliminate the cleaved adapters generated from USER II cleavage after 5’adapter ligation step, column purification was used to remove those small adapter fragments (23 and 21 nt), which may cause formation of primer-adapter dimer during the following PCR step, then affect the final yield of PCR product. In this study, we performed USER II cleavage on the ligated product and evaluated if the effect of column purification on the reduction rate of cleaved adapters is similar or higher than the test above. Our result showed that up to 91±1% of cleaved adapters were removed by column purification compared to the one without cleanup procedure after USER II cleavage process (Figure 2D, lanes 3 and 4). In brief, column purification after USER II step could clear up most cleaved adapters to obtain a clean template for the following PCR amplification, and the replacement of column purification to previous gel purification step (Figure 1A) will minimize the potential sample cross-contamination caused by gel loading and cutting, and increase the throughput of sample processing and recovery yield.

### Effect of dU Adapter on PCR amplification

To determine whether the involvement of dU adapter could increase the PCR amplification efficiency, we compared the yields of PCR product between normal adapter and dU adapter. The primary reason to design this dU adapter is to reduce the adapter-primer dimer formation during PCR process in the library preparation, and increase the efficiency in PCR amplification. Thus, 100 ng polyA RNA input was used to generate real cDNA template with normal adapter or dU adapter following related rG4-seq [23] and rG4-seq 2.0 (See methods) to perform PCR amplification experiment with increasing PCR cycles (8, 11, 14 and 17) and ran the samples on a 2% agarose gel to quantify the PCR products.

The expected PCR product (150-400 nt) were quantified and compared. Our result showed that dU adapter contributed to higher PCR efficiency than normal adapter under same PCR cycle (Figure 4). Moreover, for library preparation with 100 ng polyA-RNA input, PCR product with 11 cycles with dU adapter (Figure 2E, lane 7) was adequate for sequencing, while more than 17 PCR cycles were required with normal adapter (Figure 2E, lane 4). Unspecific PCR products from 5’dU adapter appeared at 14 and 17 cycles (Figure 2E, lanes 8 and 9), suggesting more than 14 PCR cycles could result in over-amplification using dU adapter. The overall PCR yield were significantly increased with dU adapter compared to normal adapter (Figure 2F). In conclusion, dU adapter could help to get required PCR product with less cycles than normal adapter.

**Figure 3.**
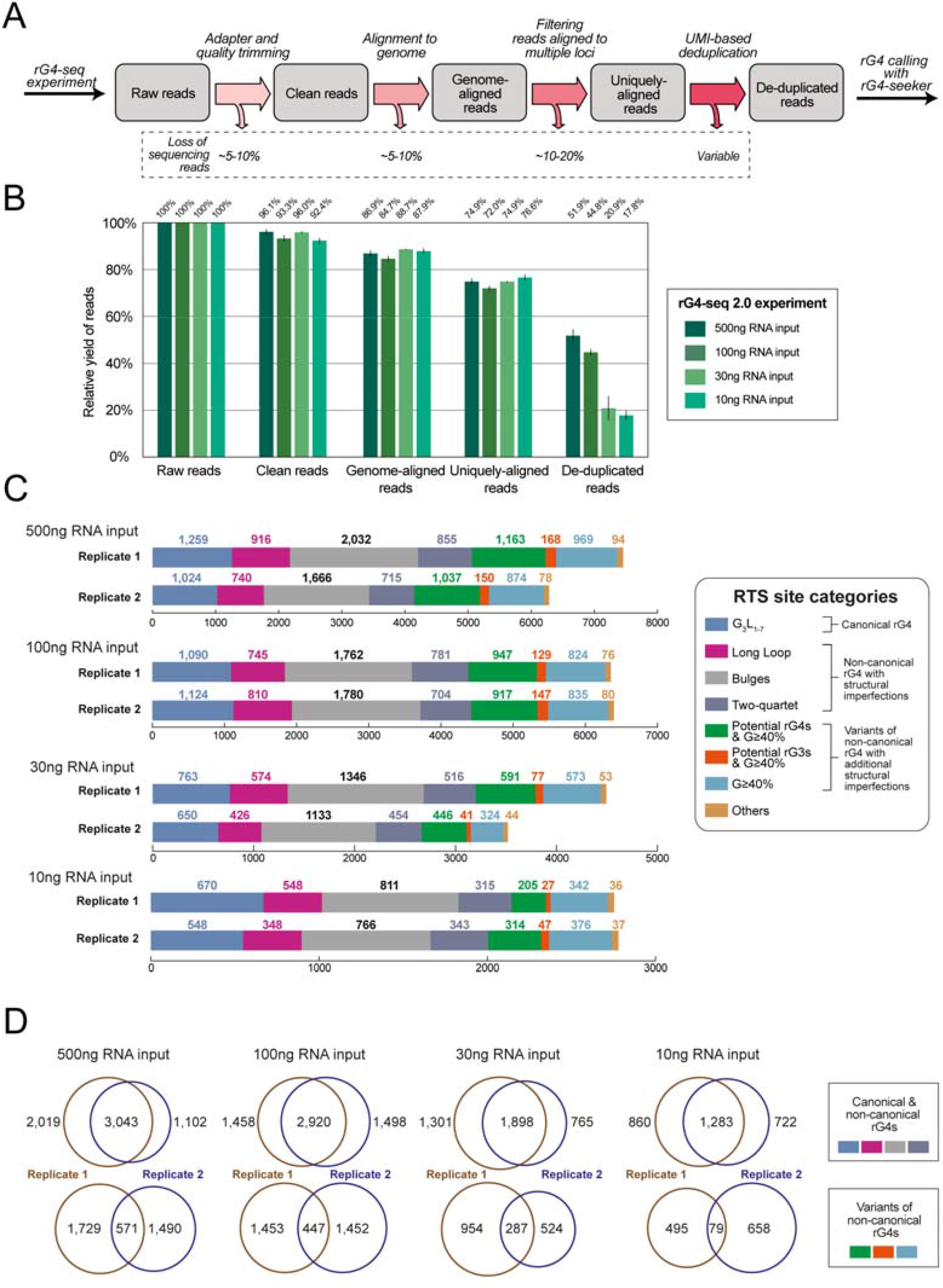
The metrics and rG4 detection outcomes of benchmarking rG4-seq 2.0 experiments. **(A)** Bioinformatic workflow of the pre-processing and deduplication of rG4-seq sequencing data. The estimated loss of sequencing reads in each step is highlighted. **(B)** Changes in the relative yield of reads in benchmarking rG4-seq 2.0 libraries through the pre-processing and deduplication steps. **(C)** Distribution of RTS sites detected in the benchmarking rG4-seq 2.0 experiments. RTS sites were assigned to rG4 structural classes according to their adjacent nucleotide sequences. RTS sites classified as “Others” were considered false positive detections as their adjacent nucleotide sequences do not satisfy the minimum requirements for forming quadruplex or triplex structures. **(D)** Reproducibility of RTS sites within biological replicates of benchmarking rG4-seq experiments.

**Figure 4.**
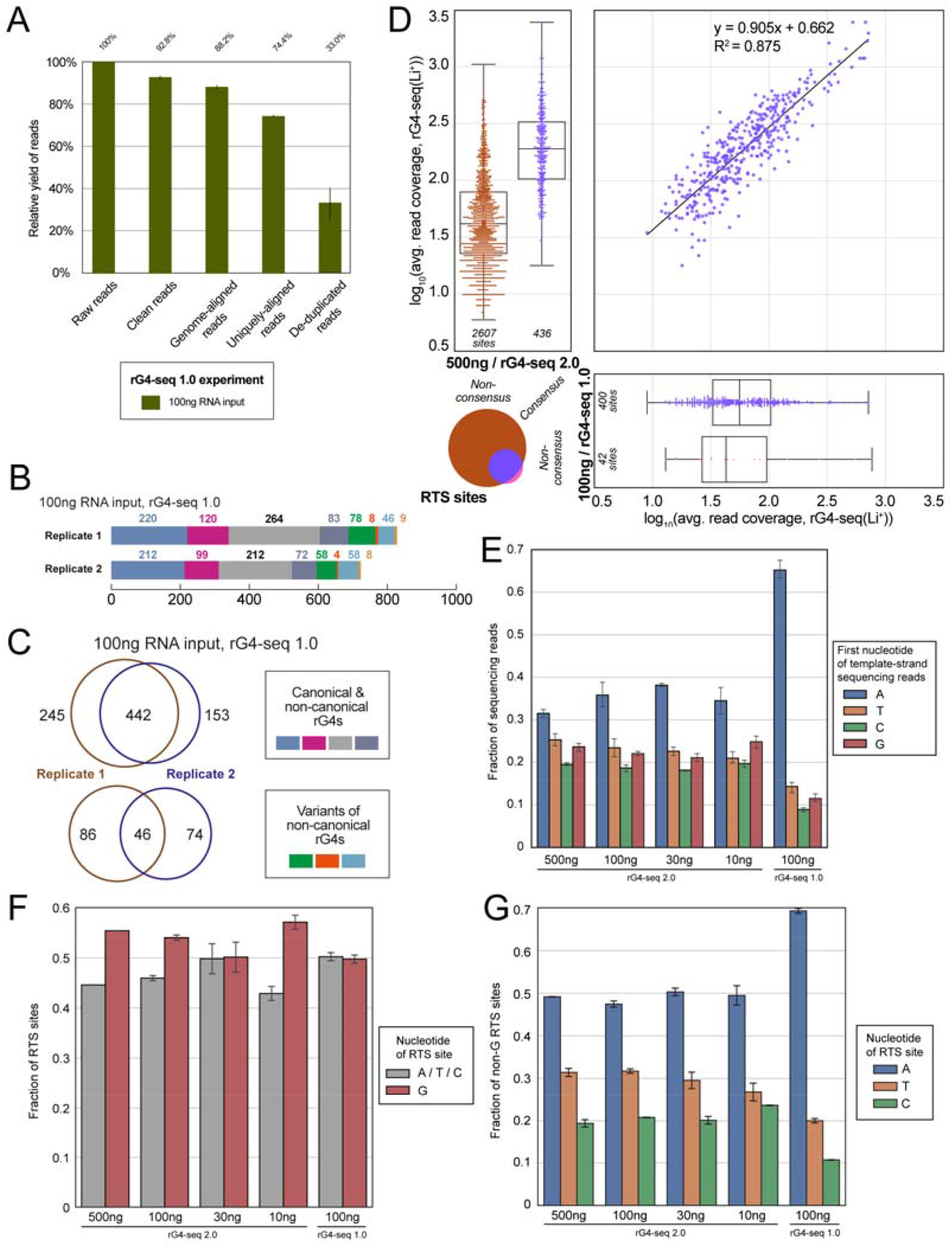
The metrics, rG4 detection outcome and underlying properties of the 100 ng-input rG4-seq 1.0 library. **(A)** Changes in the relative yield of reads in the 100 ng-input rG4-seq 1.0 library through the pre-processing and deduplication steps. **(B)** Distribution of RTS sites detected in the 100 ng-input rG4-seq 1.0 experiment. RTS sites were assigned to rG4 structural classes according to their adjacent nucleotide sequences. RTS sites classified as “Others” were considered false positive detections as their adjacent nucleotide sequences do not satisfy the minimum requirements for forming quadruplex or triplex structures. **(C)** Reproducibility of RTS sites within biological replicates of the 100 ng-input rG4-seq 1.0 experiment. **(D)** Intersection of consensus RTS sites detected in 500 ng-input rG4-seq 2.0 experiment and 100 ng-input rG4-seq 1.0 experiment. **(E)** Distribution of first nucleotides of template-strand sequencing reads in rG4-seq 2.0 and 1.0 libraries. **(F, G)** Distribution of nucleotide of RTS sites detected in rG4-seq 2.0 and 1.0 libraries, comparing **(F)** Guanine and non-Guanine RTS sites **(G)** within non-Guanine RTS sites.

### Benchmarking of rG4-seq 2.0

To benchmark the performance of the rG4-seq 2.0 protocol (Figure S5), new two-replicate rG4-seq libraries using variable inputs of HEK293T polyA-enriched RNA (500 ng, 100 ng, 30 ng and 10 ng per sample) were constructed, and sequenced with NGS at depth of approximately 100 million read-pairs per sample. The 500 ng-input library served as a baseline condition, using the RNA input amount recommended by the original rG4-seq protocol [22]. Meanwhile, the 100 ng-/30 ng-/10 ng-input libraries served to simulate rG4-seq applications with reduced RNA inputs. To evaluate the improvements introduced by 5’ dU adapter in a realistic scenario of rG4-seq application, a two-replicate rG4-seq 1.0 library using 100 ng of RNA input and normal 5’ adapter was also constructed as a control group. All rG4-seq data was then pre-processed, deduplicated using unique molecular identifiers (UMIs), and analyzed with rG4-seeker to identify rG4-induced RTS events as previously described (Figure 3A, Table S2) [24]. The workflow enabled quantitative evaluation of the influence of RNA input amount on library complexity and rG4 identification outcome.

### rG4-seq 2.0 supports rG4 identification at variable RNA input levels

Since rG4 identification with rG4-seq is established on the capturing of RTS event from unique cDNA fragments, not all sequenced reads in a library are relevant to bioinformatic analysis – only the subset of clean reads uniquely aligned to reference genome are utilized, and the UMI-based deduplication step further eliminates reads carrying duplicate information (Figure 3A). Reads discarded in different stages of analysis are considered a loss in read yield as they took up sequencing throughput without contributing to rG4 identification efforts. Our previous and proposed modification to rG4-seq protocol focused on improving read yield via reducing non-usable reads (e.g., adapter reads) and increasing the unique cDNA fragments produced (a.k.a. library complexity) per unit of RNA input.

To evaluate the yield of reads in the 4 HEK293T rG4-seq 2.0 libraries, the changes in yield through different stages of bioinformatic analysis were quantified (Figure 3B). Results suggests the change in yield from trimming, alignment and filtering for uniquely mapped reads are consistent across the 4 libraries, where 70-80% of reads were retained through these steps (Figure 3B, Table S2). The yield figures are also on-par with normal expectations of a typical RNA-seq library[43]. On the other hand, the RNA input amount affected the yield from deduplication step substantially: the baseline 500 ng-input library recorded ~30% yield loss, the 100 ng-input library recorded a ~40% loss, and the 30 ng-/10 ng-input libraries recorded ~75% loss (Figure 3B). The observation is expected as library complexity scales proportionally with input RNA amount, such that a lower input library typically would have a higher fraction of duplicated reads at the same sequencing depth. Furthermore, as the PCR duplication rates are similar between the 500 ng-input and 100 ng-input libraries despite a 5-fold difference in RNA amount, indicating the 100 million reads-pair sequencing depth may be too shallow for the complexity of the 500 ng-input library. In contrast, high duplicate rates in the 30 ng-input and 10 ng-input libraries is suggesting that a lower sequencing depth may reduce wasted throughput and improve read yield.

To evaluate the rG4 identification outcomes of individual rG4-seq 2.0 libraries, the RTS sites were first called in a replicate-independent manner and classified by their associated rG4 motifs (Figure 3C). The reproducibility of RTS sites across replicates were then evaluated (Figure 3D). Results showed that the 500 ng and 100 ng replicates captured similar number of RTS sites across different rG4 motif categories (Figure 3C), suggesting that they are equivalent in rG4 identification outcomes. Meanwhile, although the 30 ng and 10 ng replicates captured considerably less RTS sites, the relative ratio of RTS sites across motif categories remained similar (Figure 3C), suggesting both libraries would have captured a smaller, random subset of RTS sites from the HEK293T transcriptome. Also, across all libraries and replicates, the false positives (RTS sites in “Others” category) remained low at <3% (Figure 3C). Together, the result supports that rG4-induced RTS events were accurately and specifically captured using rG4-seq 2.0, regardless of input RNA amount.

On the other hand, the overlapping of RTS sites between the biological replicates were found to be ~60-70% for canonical and non-canonical rG4s (Figure 3D), where the overlapping rates matches the observations in the previous HeLa rG4-seq data [22], suggesting the replicates are well coherent with each other. While the overlapping rate of RTS sites from variant rG4 motifs were substantially lower (Figure 3D), which is not surprising as these variant rG4 motifs typically form less-stable rG4 structures with more structural imperfections, and have higher stochasticity in detection with rG4-seq. Altogether, 1,364 to 3,623 consensus (between 2 replicates) rG4 structures were detected from 10-500 ng input rG4-seq 2.0 libraries (Table S3). The union of detected rG4s totals to 5,061, among which 744 rG4s were detected across all 4 libraries (Supplementary File 1).

### 5’ dU adapter of rG4-seq 2.0 improves nucleotide bias in RTS site identification

To evaluate the effects of the new 5’ dU adapter, rG4-seq 1.0 library using 100 ng of HEK293T polyA-enriched RNA input and normal 5’ adapter was constructed and analysed. Results suggested that 100 ng-input rG4-seq 1.0 library has a yield of ~33% (Figure 4A, Table S2), which is slightly lower than ~44% of the 100 ng-input rG4-seq 2.0 library (Figure 3B, Table S2). However, significantly less rG4 motifs were detected from the rG4-seq 1.0 library (Figure 4B) in comparison to rG4-seq 2.0 libraries (Figure 3C), suggesting the performance of rG4-seq 1.0 at 100 ng RNA input is substantially subpar compared to rG4-seq 2.0. Interestingly, in contrast to the reduced rG4 detections, the reproducibility of RTS sites between replicates of the rG4-seq 1.0 library (Figure 4C) remained consistent with the observation from rG4-seq 2.0 libraries (Figure 3D). Moreover, the consensus (from two replicates, canonical and non-canonical rG4s only) set of RTS sites from the rG4-seq 1.0 library also highly overlapped with the consensus set from the 500 ng rG4-seq 2.0 library (Figure 4D). The observation suggested while most rG4s identified from the rG4-seq 1.0 library were indeed genuine, the library may have experienced an incomplete capturing of rG4-induced RTS events from the 100 ng of input RNA. Altogether, 488 consensus (between 2 replicates) rG4 structures were detected in the 100 ng-input rG4-seq 1.0 library (Table S3). The union of detected rG4s by all rG4-seq 2.0 and 1.0 libraries totals to 5,085, among which 306 rG4s were detected across all 5 libraries (Supplementary File 1).

To explore the underlying causes of the phenomenon, we first evaluated the distribution of first nucleotide in the template strand of sequencing reads after PCR deduplication (Figure 4E). These nucleotides correspond to the 5’-most nucleotides of the RNA fragments, which were ligated to the hairpin adapter (Figure 1). The rG4-seq 1.0 library was found to possess substantially A-bias in the 5’-most nucleotide (~65%); in contrast, all 4 rG4-seq 2.0 libraries showed relatively stable distributions of 5’-most nucleotides, with ~35% As and ~20% Ts, Cs and Gs (Figure 4E). The observation suggested this 5’ nucleotide bias is specific to the rG4-seq 1.0 protocol.

Since RTS information is captured by the 5’ most nucleotide of RNA fragments, we next investigated if this A-bias would have impacted the identification of rG4-induced RTS sites. It was found that consistent across all rG4-seq 1.0 and 2.0 libraries, around half of the detected RTS sites occurred on Gs, while the other half occurred on As, Ts and Cs (Figure 4F). The observation is coherent with a general property of rG4-induced RTS effects, where it typically occurs at the 3’ - most G nucleotide of the rG4 motif, and/or at the adjacent nucleotide that is not an G. However, a closer look into the distribution of non-G RTS events revealed rG4-seq 1.0 library captures substantially more RTS events on As and less on Ts/Cs, when compared with rG4-seq 2.0 libraries (Figure 4G). The results indicated that compared to rG4-seq 2.0, the rG4-seq 1.0 protocol would have introduced an intrinsic A-bias in the 5’ - most nucleotide of sequenced RNA fragments. The bias could affect the capturing of RTS sites located on non-A nucleotides, thus compromising the detection of some rG4 motifs that are flanked by Ts, Cs and Gs at its 3’ end.

Given the rG4-seq 2.0 and 1.0 libraries both shared most experimental protocols, employed 5’ adapters of identical structure (except at the cleavable dU nucleotide) and incorporated UMIs for PCR deduplication, it is unlikely that their differences in 5’-most nucleotide bias were caused by 5’ adapter ligation bias or PCR-duplicate reads. By elimination, the observation suggested the use of a 5’ dU adapter and adapter cleavage step in rG4-seq 2.0 improved the nucleotide bias in rG4-induced RTS site identification over the original hairpin adapter.

### rG4 identification performance is unaffected by a 5-fold RNA input reduction

To further characterize the influence of input RNA amounts on rG4 identification outcome, the consensus (from two replicates, canonical and non-canonical rG4s only) sets of RTS sites from rG4-seq 2.0 libraries were extracted and their intersections were calculated. Moreover, since rG4 detection outcomes are influenced by read coverage near rG4 motifs, the local coverage information of RTS sites was also incorporated in the analysis.

The intersection of RTS sites from the 500 ng-input and 100 ng-input libraries suggested that two libraries agree with each other on more than 70% of the RTS sites (Figure 5A). Importantly, the level of agreement is proportional to read coverage: nearly all RTS sites with coverage >100x are agreed, while the non-agreed sites usually have lower coverage at between 15-50x (Figure 4A). The observation is coherent with our previous findings from HeLa rG4-seq data[22], where the agreement between rG4-seq experiments is non-ideal at <32x read coverage. Nevertheless, it is important to note that low coverage RTS sites are not more prone to false positive errors, since the RTS sites shown in Figure 4A are the consensus between two biological replicates. Instead, disagreement observed should be accounted to the stochastic nature of capturing rG4-induced RTS events with relatively low number of sequencing reads. On the other hand, the read coverage of the agreed RTS sites were highly correlated between the two libraries (Figure 5A): the linear regression formula suggested a slope of ~1.0 and a small intercept of 0.132, indicating the 100 ng-input library contains similar number of reads for high-coverage RTS sites, while slightly fewer reads for low-coverage RTS sites. Together, the observation supports a conclusion that the 500 ng-input and 100 ng-input libraries are nearly equivalent in terms of rG4 identification performances.

**Figure 5.**
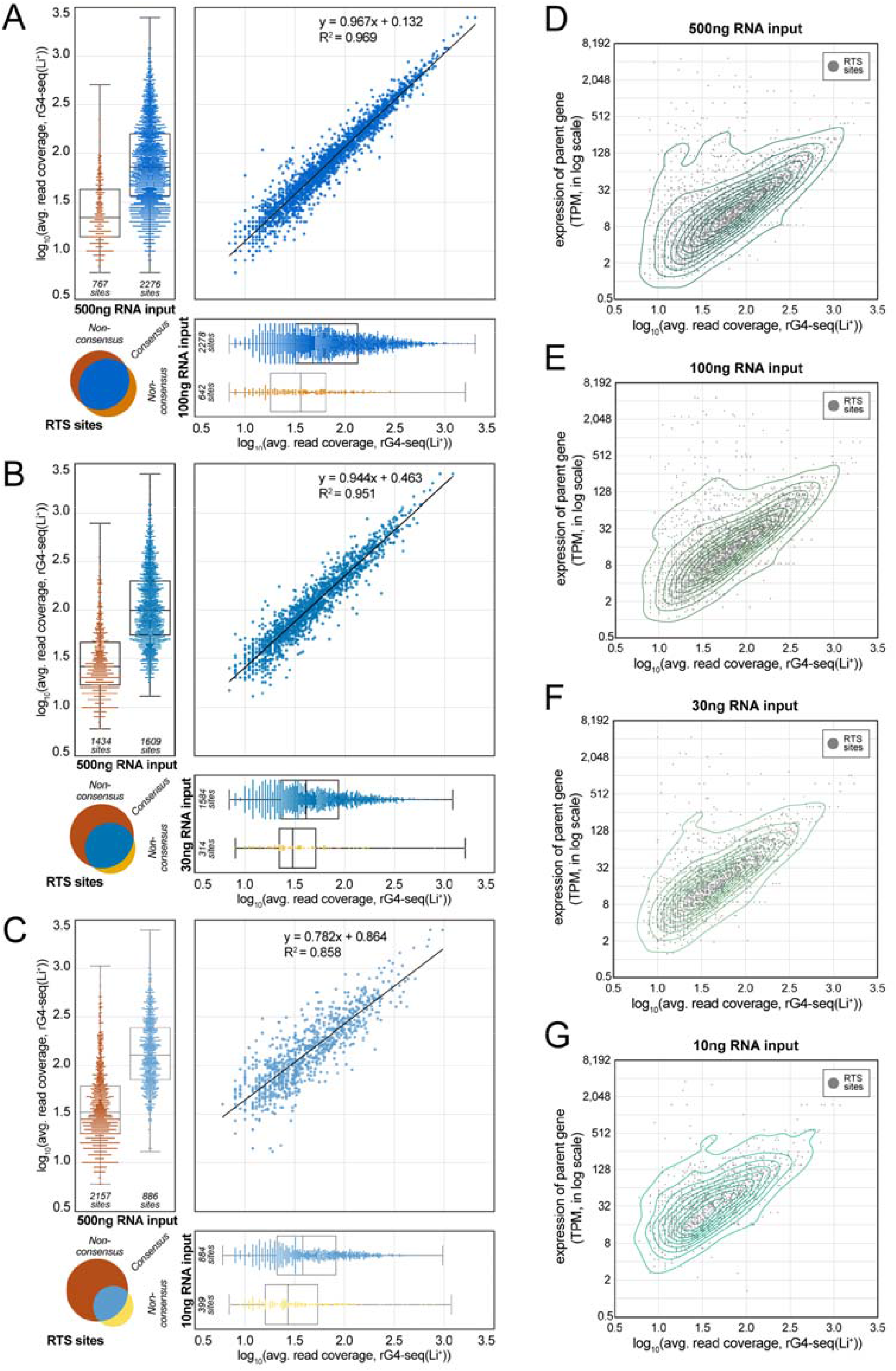
Reproducibility of RTS sites across benchmarking rG4-seq 2.0 experiments. **(A,B,C)** Intersection of consensus RTS sites detected in **(A)** 500 ng-input and 100 ng-input libraries, **(B)** 500 ng-input and 30 ng-input libraries and **(C)** 500 ng-input and 10 ng-input libraries. The Venn diagrams indicate the number of RTS sites that are agreed or non-agreed between the two libraries. The swarm plots and overlayed box plots display and contrast the differences in read coverages of agreed/non-agreed RTS sites. The scatter plots indicate the correlation and linear regression outcomes of read coverage of the RTS sites agreed between two rG4-seq libraries. **(D, E, F, G)** Average read coverages and relative abundances of the parent gene of the consensus RTS sites detected in the rG4-seq experiments. The contour lines from kernel density estimation were overlayed to help illustrating the density distribution of RTS sites.

### Further RNA input reduction impacts the detection of low coverage RTS sites

On the other hand, the 30 ng-input and 10 ng-input libraries produced different results in the intersection analysis. From the observation that respectively ~85%/~70% of RTS sites detected in the 30 ng-/10 ng-input library agree with the sites detected in the 500 ng-input library (Figure 5B, C), it can be confirmed that rG4 discoveries from the two libraries are genuine.

However, a correlation analysis indicated that consensus RTS sites in these two libraries had substantially lower read coverage than their matched counterparts in the 500 ng-input library (Figure 5B, C) – around 2x less coverage for the 30 ng-input library sites, and 3-5x less for the 10 ng-input library sites. Moreover, in the 30 ng-input library, the linear regression formula has a slope of ~0.8, indicating the undesirable reduction in coverage tends to be amplified in the already-low coverage regions of the transcriptome. Importantly, the observation explains why the 30 ng-/10 ng-input library had fewer RTS detection and was unable to rediscover many of RTS sites in the 500 ng-input library: the majority of the non-rediscovered RTS sites have a read coverage of 15-50x in the 500 ng-input library, and the projected read coverage of these sites in a 30 ng-/10 ng-input scenario are either at near or at below the detection limit of rG4-seq experiments (at around 16x coverage). Together, our findings suggested that while accurate rG4 identification with rG4-seq 2.0 is possible at sub-30 ng levels of RNA input, the sensitivity for rG4s in low coverage regions of the transcriptome might be substantially affected.

### A reference guideline for RNA input level options with rG4-seq 2.0

Finally, to outline the relationships between transcript abundances and the sensitivity for rG4s in a rG4-seq 2.0 experiment, we plotted the read coverage of the consensus RTS sites against the abundances (in TPM, transcript per million) of their parent genes for the 4 libraries (Figure 5D, E, F, G). The plots corresponding to the 500 ng-input (Figure 5D) and 100 ng-input library (Figure 4E) showed that the majority of RTS site were detected from genes of TPM 0.5 - 16, and the local read coverage for these RTS site ranged between 10x to 100x. In contrast, owing to the general reduction in read coverage, RTS detections from these lower abundances genes was impacted in the 30 ng-input library (Figure 5F) while largely compromised in the 10 ng-input library (Figure 5G).

Together, our benchmarking of rG4-seq 2.0 suggested the following conclusions:

1. Regardless of input RNA amount, rG4-induced RTS events can be accurately captured to reveal the presence of rG4 structures in HEK293T transcriptome.
2. Given a 100 million read-pair sequencing depth, a 100 ng poly-A enriched RNA input per sample would be optimal using the new protocol. A higher input RNA amount would require higher sequencing depth to fully capture the increased library complexity.
3. Using an RNA input amount below 100 ng would reduce usable read coverage in lower abundance genes, increase the stochasticity in rG4 detection from these genes, and has risk of compromising rG4 identification if the coverage falls below the detection limit.

In practice, if a comprehensive genome-wide profiling of rG4s is desired, users are advised to use 100 ng-input RNA/100 million read pairs as a reference point and scale the RNA amount and sequencing throughput proportionally for optimal read yield. A very low RNA input at 10 ng levels would only be desirable if the required scope of rG4 detection is limited to higher abundance genes, typically at >10 TPM. Meanwhile, if the detection of rG4s from rare transcripts (e.g. long non-coding RNAs) via coupling target-capture methods with rG4-seq 2.0 is desired, a practical guideline could be:

1. Relative abundances of targeted genes should be ≥4 TPM post-enrichment.
2. If the abundances of targeted genes post-enrichment is in range 4-8 TPM, at least 100 ng of post-enrichment RNA should be used
3. If the abundances of targeted genes post-enrichment is in range 8-16 TPM, at least 30 ng of post-enrichment RNA should be used.
4. If the abundances of targeted genes post-enrichment is in range ≥16 TPM, at least 10 ng of post-enrichment RNA should be used.

## CONCLUSIONS

In summary, we developed the rG4-seq 2.0 protocol by incorporating the use of dU adapter, eliminating the gel purification steps and optimizing the experimental procedures, yielding a more user-friendly and efficient pipeline than the original rG4-seq method. We firstly applied the rG4-seq 2.0 to purified polyA-enriched RNA from HEK293T cell line and generated 4 cDNA rG4-seq libraries using decreased polyA-RNA inputs (500 ng, 100 ng, 30 ng and 10 ng). All libraries showed good qualities and reproducibility with high mapping rate of transcript. rG4-induced RTS events can be accurately captured by rG4-seq 2.0 to reveal the presence of rG4 structures in HEK293T transcriptome. Moreover, the new dU adapter in rG4-seq 2.0 improved rG4 calling outcome by mitigating a significant nucleotide bias in rG4-induced RTS site identification persistent in the old method. Last, we outlined the relationships between transcript abundances and the sensitivity for rG4s in human transcriptome and proposed a standard recommendation, which provides the amount of RNA inputs and sequencing depth for specific rG4 study based on the target abundance in the transcriptome. The benchmarking on rG4-seq 2.0 will provide a new and important platform for constructing cDNA library for wide-ranging biological applications. It will not only be applicable in rG4-seq and RNA structurome studies, but also to other RNA-related next generation sequencing applications such as RNA-seq.

## DATA AVAILABILITY

The data generated during all experiments is available for public. Sequencing datasets generated in this study are available in the NCBI SRA repository under the accession number PRJNA728962.

(https://dataview.ncbi.nlm.nih.gov/object/PRJNA728962?reviewer=l2nlutg0mjolj4crba2v3pstq7)

## SUPPLEMENTARY DATA

Supplementary Data are available at RNA online.

## ACKNOWLEDGEMENT

We thank members from the Kwok Lab and Chan Lab for the discussion and support. Any opinions, findings, conclusions or recommendations expressed in this publication do not reflect the views of the Government of the Hong Kong Special Administrative Region or the Innovation and Technology Commission.

