## Supporting Information for "rG4-seq 2.0: enhanced transcriptome-wide RNA G-quadruplex structure sequencing for low RNA input samples"

**Table of Contents**

**Table S1.** DNA oligonucleotides used in this study

**Table S2.** Number of usable sequencing reads in respective bioinformatic analysis steps

**Table S3.** Number of rG4 motifs identified in rG4-seq libraries

**Figure S1.** Effect of different conditions on the 5' dU adapter cleavage

**Figure S2.** Effect of uracil position on adapter cleavage.

**Figure S3.** The effect of dU adapter concentration on ligation efficiency

**Figure S4.** Effect of column purification on the 106-nt mimic ligated product

**Figure S5.** Experimental flowchart of RNA G-quadruplex structure sequencing 2.0 (rG4-seq 2.0)

**Supplementary File 1 (uploaded separately).** Details of identified rG4 in each rG4-seq library (in xlsx format) and the lists of consensus rG4s / union rG4s across rG4-seq libraries (in BED format)

**Table S1.** DNA oligonucleotides used in this study

| Name of Oligo | Reaction | Length (nt) | Sequence | Modification |
|---|---|---|---|---|
| Normal 5' adapter | dU Cleavage | 44 | 5'-/5Phos/AGATCGGAAGAGCGTCGTGTAGCTCTTCCGATCTN$_{10}$/3SpC3/-3' | 5'-P; 3'-SpC3 |
| 5' dU Adapter | dU Cleavage | 44 | 5'-/5Phos/AGATCGGAAGAGCGTCGTGTAGC/dU/CTTCCGATCTN$_{10}$/3SpC3/-3' | 5'-P; 3'-SpC3 |
| dU Loop18 adapter | dU Cleavage | 44 | 5'-/5Phos/AGATCGGAAGAGCGTCG/dU/GTAGCTCTTCCGATCTN$_{10}$/3SpC3/-3' | 5'-P; 3'-SpC3 |
| dU Loop20 adapter | dU Cleavage | 44 | 5'-/5Phos/AGATCGGAAGAGCGTCGTG/dU/AGCTCTTCCGATCTN$_{10}$/3SpC3/-3' | 5'-P; 3'-SpC3 |
| dU Stem27 adapter | dU Cleavage | 44 | 5'-/5Phos/AGATCGGAAGAGCGTCGTGTAGCTCT/dU/CCGATCTN$_{10}$/3SpC3/-3' | 5'-P; 3'-SpC3 |
| dU Stem32 adapter | dU Cleavage | 44 | 5'-/5Phos/AGATCGGAAGAGCGTCGTGTAGCTCTTCCGA/dU/CTN$_{10}$/3SpC3/-3' | 5'-P; 3'-SpC3 |
| dU Stem34 adapter | dU Cleavage | 44 | 5'-/5Phos/AGATCGGAAGAGCGTCGTGTAGCTCTTCCGATC/dU/N$_{10}$/3SpC3/-3' | 5'-P; 3'-SpC3 |
| N40+22nt DNA | 5' Adapter Ligation | 62 | 5'-N$_{40}$TCTAGCCTTCTCGTGTGCAGAC-3' | 5'-OH; 3'-OH |

| | | | | |
|---|---|---|---|---|
| 106-nt mimic ligated product | Column purification recovery | 106 | 5'-CAGACGTGTGCTCTTCCGATCTNNNNNNNNNNNNNNNNNNNNNNNNNNNNNNNNNNNNNNNAGATCGGAAGAGCGTCGTGTAGCTCTTCCGATCTNNNNNNNNNNN-3' | 5'-OH; 3'-OH |
| 3'-Adapter | 3'-RNA Ligation | 28 | 5'-/5rApp/NNNNNNAGATCGGAAGAGCACACGTCTG/3SpC3/-3' | 5'-rApp; 3'-SpC3 |
| Reverse primer | Reverse transcription | 22 | 5'-CAGACGTGTGCTCTTCCGATCT-3' | 5'-OH; 3'-OH |
| Forward Primer | PCR Amplification | 58 | 5'-AATGATACGGCGACCACCGAGATCTACACTCTTTCCCTACACGACGCTCTTCCGATCT-3' | 5'-OH; 3'-OH |
| Reverse Primer | PCR Amplification | 58+6 | 5'-CAAGCAGAAGACGGCATACGAGAT-(6 nt index_seq)-GTGACTGGAGTTCAGACGTGTGCTCTTCCGATCT-3' | 5'-OH; 3'-OH |

**Table S2.** Number of usable sequencing reads in respective bioinformatic analysis steps

| **rG4-seq 2.0 Library** | **Raw read pairs** | **Clean read pairs** | **Mapped read pairs** | **Uniquely-aligned read pairs** | **Deduplicated read pairs** |
|---|---|---|---|---|---|
| 500ng-K-rep1 | 137,398,397 | 133,167,732 | 120,159,372 | 103,560,827 | 68,125,003 |
| 500ng-K-rep2 | 88,985,820 | 84,838,879 | 77,011,859 | 66,541,953 | 49,331,989 |
| 500ng-Li-rep1 | 166,406,649 | 162,151,063 | 147,114,022 | 127,195,204 | 82,133,712 |
| 500ng-Li-rep2 | 100,566,437 | 95,288,401 | 85,623,158 | 73,466,003 | 53,703,838 |
| 100ng-K-rep1 | 110,410,923 | 103,333,453 | 93,679,546 | 79,544,932 | 48,137,468 |
| 100ng-K-rep2 | 122,039,081 | 112,433,798 | 102,468,376 | 87,309,462 | 57,260,162 |
| 100ng-Li-rep1 | 97,102,071 | 92,672,652 | 83,919,534 | 71,310,823 | 42,817,726 |
| 100ng-Li-rep2 | 126,122,160 | 116,111,852 | 105,244,652 | 89,419,592 | 56,094,552 |
| 30ng-K-rep1 | 93,804,399 | 89,440,824 | 83,022,991 | 70,261,384 | 26,488,259 |
| 30ng-K-rep2 | 165,418,442 | 158,976,261 | 146,362,570 | 123,440,346 | 23,378,203 |
| 30ng-Li-rep1 | 121,866,093 | 117,261,135 | 108,314,277 | 91,499,372 | 23,861,539 |
| 30ng-Li-rep2 | 109,402,650 | 105,156,448 | 97,080,342 | 81,955,634 | 23,645,790 |
| 10ng-K-rep1 | 114,115,611 | 104,060,606 | 98,905,474 | 86,175,293 | 23,158,515 |
| 10ng-K-rep2 | 94,216,637 | 88,586,738 | 84,416,825 | 74,012,659 | 16,809,716 |
| 10ng-Li-rep1 | 124,962,015 | 115,322,248 | 110,028,002 | 95,541,621 | 22,152,141 |
| 10ng-Li-rep2 | 74,104,271 | 68,206,790 | 64,690,588 | 56,222,720 | 11,346,437 |
| **rG4-seq 1.0 Library** | **Raw read pairs** | **Clean read pairs** | **Mapped read pairs** | **Uniquely-aligned read pairs** | **Deduplicated read pairs** |
| 100ng-K-rep1 | 145,927,820 | 134,863,585 | 128757741 | 107,794,352 | 36,205,284 |
| 100ng-K-rep2 | 130,054,116 | 120,094,560 | 113843448 | 96,142,554 | 44,390,570 |
| 100ng-Li-rep1 | 83,877,386 | 78,619,804 | 74856135 | 62,552,149 | 37,302,028 |
| 100ng-Li-rep2 | 115,133,457 | 106,623,017 | 100956422 | 86,458,904 | 32,975,862 |

**Table S3.** Number of rG4 motifs identified in rG4-seq libraries

| | HEK293-rG4-seq-2.0-10ng | | | HEK293-rG4-seq-2.0-30ng | | | HEK293-rG4-seq-2.0-100ng | | | HEK293-rG4-seq-2.0-500ng | | | HEK293-rG4-seq-1.0-100ng | | |
|---|---|---|---|---|---|---|---|---|---|---|---|---|---|---|---|
| ***rG4 motifs*** | **K-rep1** | **K-rep2** | **Con-sensus** | **K-rep1** | **K-rep2** | **Con-sensus** | **K-rep1** | **K-rep2** | **Con-sensus** | **K-rep1** | **K-rep2** | **Con-sensus** | **K-rep1** | **K-rep2** | **Con-sensus** |
| *canonical/G3L1-7* | 670 | 548 | 449 | 763 | 650 | 531 | 1090 | 1124 | 853 | 1259 | 1024 | 883 | 220 | 212 | 165 |
| *longloop* | 347 | 348 | 229 | 574 | 426 | 343 | 745 | 810 | 560 | 916 | 740 | 612 | 120 | 99 | 74 |
| *bulges* | 811 | 766 | 453 | 1346 | 1133 | 793 | 1762 | 1780 | 1146 | 2032 | 1666 | 1180 | 264 | 212 | 157 |
| *two-quartet* | 315 | 343 | 152 | 516 | 454 | 231 | 781 | 704 | 361 | 855 | 715 | 368 | 83 | 72 | 46 |
| ***subtotal : canonical & non-canonical motifs*** | ***2143*** | ***2005*** | ***1283*** | ***3199*** | ***2663*** | ***1898*** | ***4378*** | ***4418*** | ***2920*** | ***5062*** | ***4145*** | ***3043*** | ***687*** | ***595*** | ***442*** |
| *potential G-quadruplex & G>=40%* | 205 | 314 | 59 | 591 | 446 | 233 | 947 | 917 | 364 | 1163 | 1037 | 431 | 78 | 58 | 37 |
| *potential G-triplex & G>=40%* | 27 | 47 | 3 | 77 | 41 | 20 | 129 | 147 | 27 | 168 | 150 | 34 | 8 | 4 | 3 |
| *G>=40%* | 342 | 376 | 17 | 573 | 324 | 34 | 824 | 835 | 56 | 969 | 874 | 106 | 46 | 58 | 6 |
| ***subtotal : variants of non-canonical motifs*** | ***574*** | ***737*** | ***79*** | ***1241*** | ***811*** | ***287*** | ***1900*** | ***1899*** | ***447*** | ***2300*** | ***2061*** | ***571*** | ***132*** | ***120*** | ***46*** |
| ***Total – all rG4 motifs*** | ***2753*** | ***2779*** | ***1364*** | ***4440*** | ***3474*** | ***2185*** | ***6354*** | ***6397*** | ***3370*** | ***7456*** | ***6284*** | ***3623*** | ***819*** | ***715*** | ***488*** |
| *unknown* | 36 | 37 | 2 | 53 | 44 | 2 | 76 | 80 | 3 | 94 | 78 | 9 | 9 | 8 | 1 |

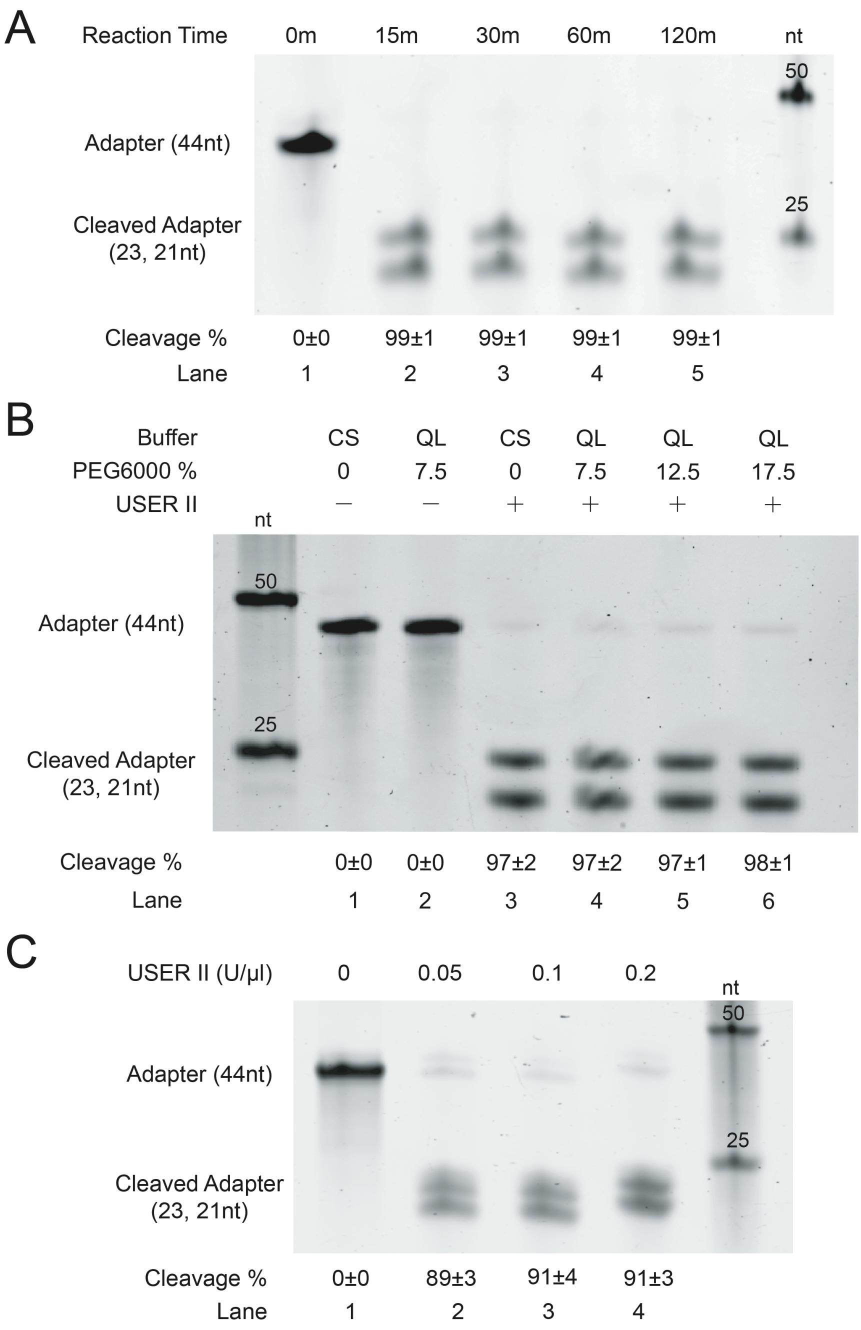

**Figure S1.** Effect of different conditions on the 5' dU adapter cleavage. **(A)** The effect of reaction time over 15min on 5' dU adapter cleavage. The figure shows the cleavage of dU adapter (lanes 2-5) with USER II at 0, 15, 30, 60 and 120 min. There is no cleavage observed at 0 min (lane 1). The cleavage rate reaches over 99% in all reaction time from 15 minutes to 2 hours (lanes 2-5). **(B)** The effect of reaction buffer and different percentage of PEG 6000 on the 5' dU adapter cleavage. The figure shows the dU cleavage under CutSmart (CS) buffer or Quick Ligase (QL) Buffer (Lanes 3-6) and with the percentage of PEG 6000 ranging from 7.5%-17.5% (Lanes 4-6). No cleavage is observed without USER II addition regardless reaction buffer type (Lanes 1-2). The cleavage rates are 97±2% under both CS buffer and QL buffer (Lane 3 and 4). From 7.5%-17.5% PEG6000 concentration under QL buffer, the % of adapter cleavage are 97-98% (Lanes 4-6), which are similar with the one under CS buffer (Lane 3). **(C)** Effect of enzyme concentration on 5’ dU adapter cleavage. Different enzyme concentrations from 0-0.2 U/µl are tested to evaluate the dU adapter cleavage rate. It reaches highest efficiency with 0.1 U/µl (Lane 3). Higher USER II concentration could not increase the cleavage rate (Lane 4). The DNA size marker indicates the fragment size. Equation 1 (See Methods) is used for calculation. The errors showed are standard deviation. nt=nucleotide; n=3.

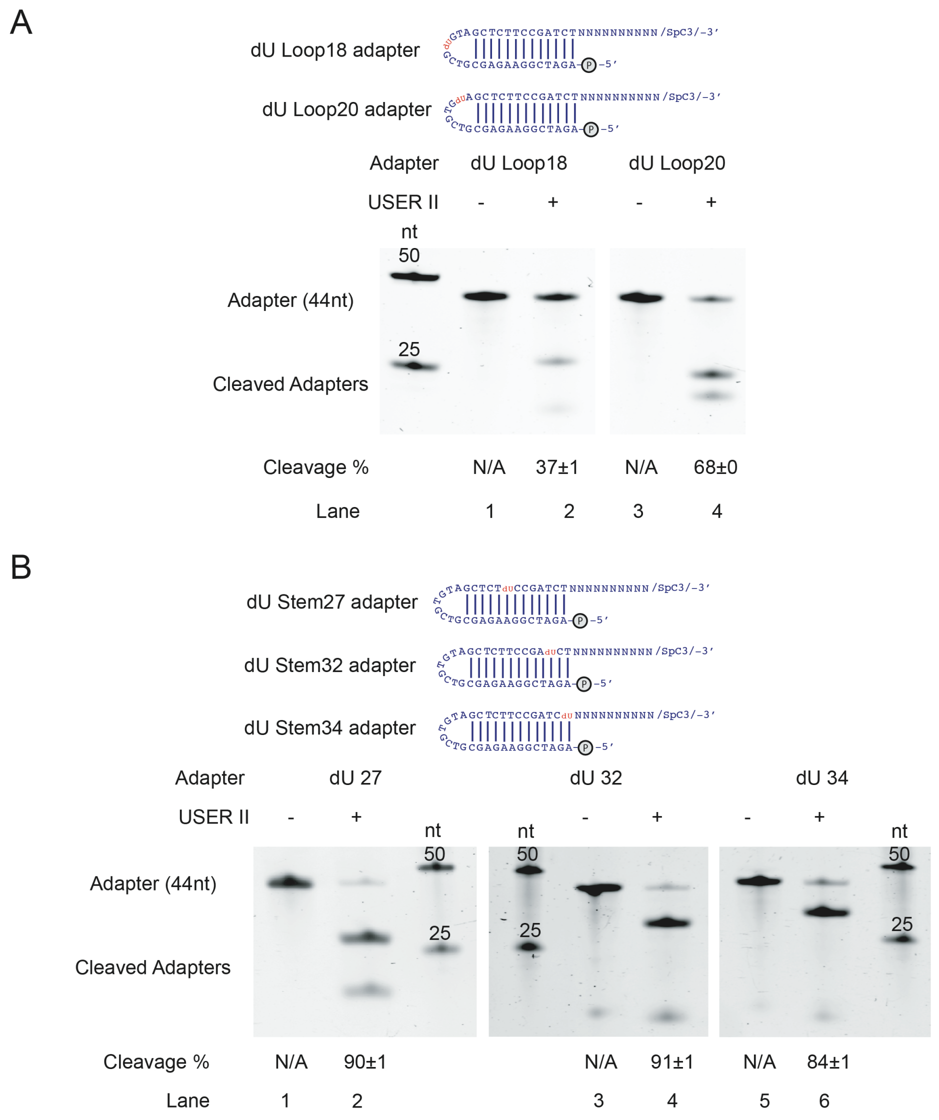

**Figure S2.** Effect of uracil position on adapter cleavage. **(A)** dU cleavage on dU loop adapters. No cleavage is observed on the dU Loop18 adapter and dU Loop20 adapter without USER II treatment (Lanes 1 and 3). the cleavage rate of dU Loop18 adapter is 37±1% (Lane 2), and for dU Loop20 adapter, the cleavage rate is 68±1% (Lane 4), which showed lower cleavage efficacy than 5' dU adapter (Figure 2, lane 7). **(B)** dU cleavage on dU stem adapters. The 3 stem adapters without USER II treatment were used as negative controls (Lanes 1, 3 and 5). The dU cleavage rates are 90±1%, 91±1% and 84±1% for Stem27 adapter, dU Stem32 adapter and dU Stem34 adapter (Lanes 2, 4 and 6), individually, which are much higher than that of dU Loop adapters (Lanes 2 and 4) showed in (A). The DNA size marker indicates the fragment size. Equation 1 (See Methods) is used for calculation. The errors showed are standard deviation. nt=nucleotide; n=3.

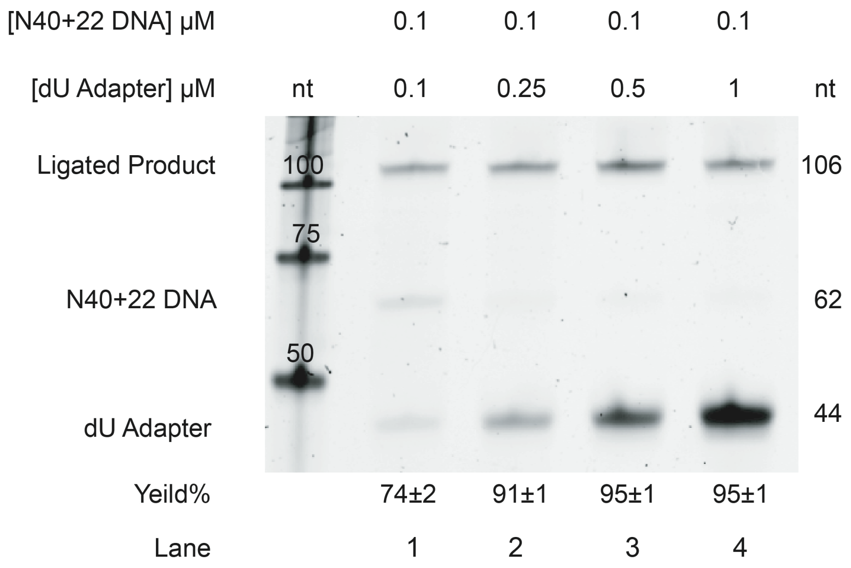

**Figure S3.** The effect of dU adapter concentration on ligation efficiency. The 106-nt ligated product is generated with varying cDNA: adapter ratio, from 1:1 to 1:10. The ligation yield enhanced with the decreasing DNA: adapter ratio and it reaches to the highest in 1:5 and 1:10 of DNA:adapter ratios (Lanes 3 and 4). Equation 2 is used for the yield% calculation (see Methods). The errors showed are standard deviation. nt=nucleotide; n=3.

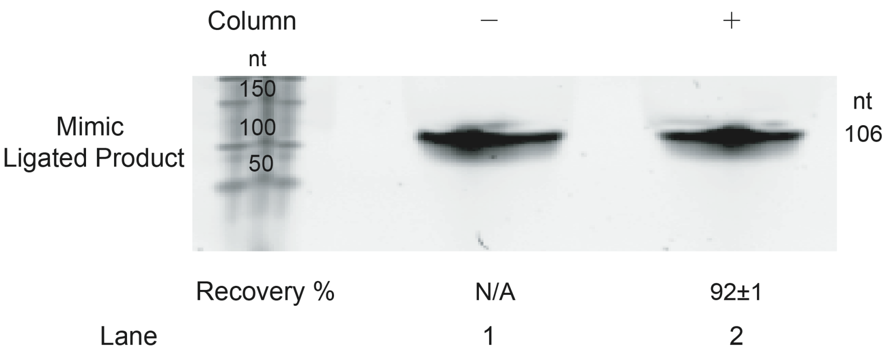

**Figure S4.** Effect of column purification on the 106-nt mimic ligated product. The column recovery rate on the ligated product is 92±1% (Lane 2). The band of 106-nt mimic ligated product without column purification (Lane 1) is defined as reference to calculate the recovery rate. Equation 4 is used for the recovery rate of calculation (see Methods). The error showed are standard deviation. nt=nucleotide; n=3.

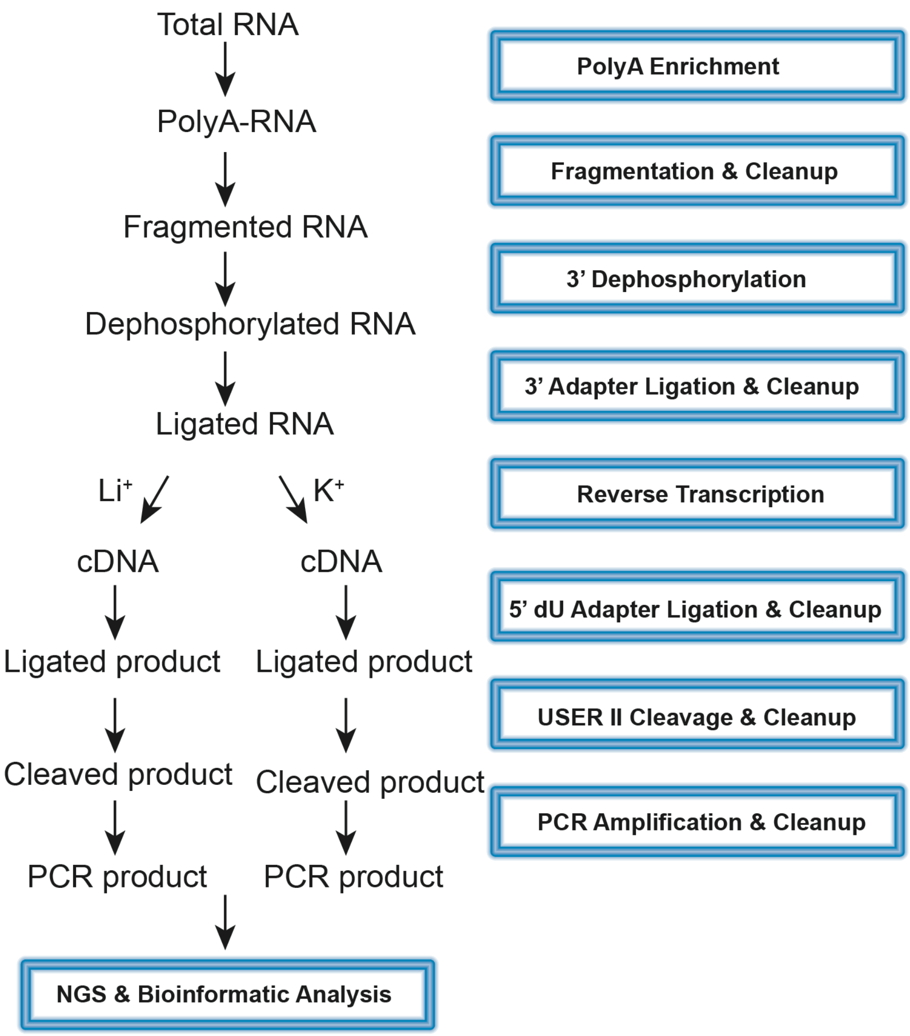

**Figure S5.** Experimental flowchart of RNA G-quadruplex structure sequencing 2.0 (rG4-seq 2.0). PolyA-enriched RNA (polyA-RNA) was obtained after 2 rounds of enrichment from total RNA. Around 250 nt random RNA fragments were generated during fragmentation step. Dephosphorylation reaction replaces the 3′-P of RNA with 3′-OH for the following 3' adapter ligation. 3′ adapter is then ligated to RNA fragment followed by excess adapter digestion and removal. Reverse transcription (RT) is performed under either $Li^+$ or $K^+$ condition. After the RT, 5′dU adapter is then ligated to cDNA followed by ligation cleanup. USER II enzyme cleaves the dU in the ligated product followed by the excess adapters cleanup. PCR amplification is performed afterwards to produce DNA library. The purified cDNA library is then sent for next-generation sequencing (NGS) followed by bioinformatic analysis.
